# MrHAMER yields highly accurate single molecule viral sequences enabling analysis of intra-host evolution

**DOI:** 10.1101/2021.01.27.428469

**Authors:** CM Gallardo, S Wang, DJ Montiel-Garcia, SJ Little, DM Smith, AL Routh, BE Torbett

**Affiliations:** Department of Immunology and Microbiology, The Scripps Research Institute, La Jolla, CA; Center for Immunity and Immunotherapies, Seattle Children’s Research Institute, Seattle, WA; Department of Integrative Structural and Computational Biology, The Scripps Research Institute, La Jolla, CA; Division of Infectious Diseases and Global Public Health, University of California, San Diego, La Jolla, CA; Veterans Affairs Healthcare System, San Diego, CA, USA; Department of Biochemistry and Molecular Biology, University of Texas Medical Branch, Galveston, TX; Sealy Center for Structural Biology, University of Texas Medical Branch, Galveston, TX; Department of Pediatrics, University of Washington School of Medicine, Seattle, WA

## Abstract

Technical challenges remain in the sequencing of RNA viruses due to their high intra-host diversity. This bottleneck is particularly pronounced when interrogating long-range co-evolution given the read-length limitations of next-generation sequencing platforms. This has hampered the direct observation of long-range genetic interactions that code for protein-protein interfaces with relevance in both drug and vaccine development. Here we overcome these technical limitations by developing a nanopore-based long-range viral sequencing pipeline that yields accurate single molecule sequences of circulating virions from clinical samples. We demonstrate its utility in observing the evolution of individual HIV Gag-Pol genomes in response to antiviral pressure. Our pipeline, called Multi-read Hairpin Mediated Error-correction Reaction (MrHAMER), yields >1000s viral genomes per sample at 99.9% accuracy, maintains the original proportion of sequenced virions present in a complex mixture, and allows the detection of rare viral genomes with their associated mutations present at <1% frequency. This method facilitates scalable investigation of genetic correlates of resistance to both antiviral therapy and immune pressure, and enable the identification of novel host-viral and viral-viral interfaces that can be modulated for therapeutic benefit.

## Introduction

Advances in viral genomics have provided new understandings of the genetic changes that underlie the pathogenesis, evolution, and epidemiology of emerging viruses. Currently, viral genomics informs the detection and management of drug resistance to antiviral therapy in rapidly mutating viruses such as HIV, Influenza[1], HCV[2], and HBV[3]. Recently, the development of distributed viral genomics methodologies have allowed successful epidemiological surveillance of SARS-CoV2, ebolaviruses, Zika virus, and other emerging viruses [4, 5]. However, technical challenges remain in the sequencing of rapidly-mutating RNA viruses which, due to their high intra-host diversity, require specialized sequencing approaches to allow the unambiguous identification of mutation patterns[6]. This bottleneck is particularly pronounced when interrogating long-range co-evolution given read-length limitations of next-generation sequencing (NGS) platforms, which hamper the direct observation of long-range genetic interactions that code for protein-protein interfaces with relevance in both drug and vaccine development [7, 8].

HIV serves as an ideal model for investigating the genetic mechanisms of RNA viruses that lead to increased fitness since it has *both* a high mutation rate and high recombination potential which, coupled with its proliferative capacity, result in prodigious viral sequence diversity in circulating virions [9, 10]. The resulting viral ‘swarm’ is highly dynamic and subject to immune and antiviral selection pressure, which can lead to accumulation of low frequency mutations that can interact cooperatively at the functional level and confer phenotypes that undercut host defenses and therapeutic interventions [11]. During combination antiretroviral therapy (cART), viral species containing drug resistance mutations (DRMs) can gradually become enriched and result in drug failure if they become the dominant species. Although the selection of DRMs is highly constrained by the need to preserve RNA secondary structure and structural stability of proteins, and to retain function of viral-viral interfaces [12, 13], numerous DRMs have been detected and catalogued in several databases (HIV LANL, Stanford HIVDB), with some DRMs found to be preferentially associated with single-nucleotide variants (SNVs) present at another loci. This process, whereby one mutation’s impact on fitness is dependent on the interactions between genes at different loci is termed functional epistasis, and has been shown to operate in HIV drug-resistance [14-16]. However, the short read-lengths of NGS prevents the direct detection of long-range pairs including correlated mutations between Gag and Pol Polyproteins and between other regions throughout the 9.2kb HIV genome. This has limited analysis of long-range interactions to bioinformatic or computational inference [7, 17], which has been shown to be unreliable given the intra-host genetic diversity in HIV [18]. Thus, the importance of identifying long-range pairs (or higher order patterns) of residue substitution in the HIV genome, and other RNA viruses is an outstanding question in the field that has been limited by technological hurdles.

Multiple approaches have been used to address current read-length limitations in viral sequencing. Single-molecule sequencing technologies can generate very long sequence reads, but due to higher single-pass error rates of 5-10% (vs. <0.1% for Illumina) minor viral variants in the population might not be accurately identified [19]. The current error rates of these long-read platforms has required multiple passes on a single molecule in order to obtain more accurate reads (i.e. HiFi reads for PacBio or 1D^2^ in ONT) [20]. Another approach is adopting novel library preparation techniques that allow reconstruction of full-length, single-molecule genomes using overlapping barcoded short reads. Three recent approaches using overlapping barcoded reads have been reported in a viral sequencing context, all relying on molecular barcoding and complex library preparation steps [21-23]. These provide accurate data and effectively longer read lengths, albeit with complex bioinformatic pipelines. However, the lack of validation of RT-PCR conditions that minimize artifactual recombination and sampling bias during amplification of viral cDNA remains an unavoidable challenge [24] that is not actively addressed in these approaches. This is critical for accurate characterization of long-range linkage, since extensive PCR generates chimeras that can either erase evidence of long-range linkage and/or generate false-positives of long-range interactions at rates of up to 30% [25]. The limiting amounts of viral RNA present in clinical isolates add a challenge as these samples generally require extensive amplification by PCR or similar isothermal methods. This requires optimization of PCR conditions to ensure sufficient yield for sequencing while maintaining accurate long-range linkage.

To address these shortcomings, we have developed a nanopore-based long-range sequencing pipeline that yields accurate single molecule sequences of circulating virions from clinical samples. We dubbed this approach Multi-read Hairpin Mediated Error-Correction Reaction (MrHAMER), and demonstrate its utility in observing the evolution of individual HIV Gag-Pol genomes in response to antiviral pressure. MrHAMER is facilitated by a fully-validated RT-PCR pipeline that amplifies cDNA from clinical isolates with minimal viral load, while preserving long-range linkage and minimizing template switching artifacts. Amplified cDNA is then sequenced to high accuracy using a template preparation and bioinformatic pipeline that generates sequential sense and antisense repeats of a cDNA molecule to reduce the inherent error rate of ONT sequencing by two orders of magnitude (to 99.9% accuracy). The MrHAMER pipeline is designed to leverage the Oxford Nanopore Technologies (ONT) MinION platform which allows for portable on-demand sequencing at a low upfront capital investment and short turnaround times, and facilitates more distributed clinical virology workflows and field deployment. We show that MrHAMER yields >1000s genomes per sample, maintains the original proportion of linked mutations present in a complex mixture, and allows the detection of single RNA genomes (and associated linked mutations) present at <1% frequency. We demonstrate and validate the utility of MrHAMER by sequencing longitudinal HIV isolates from a patient undergoing virological failure, and show a hard selective sweep [26, 27] of a pre-existing Gag-Pol genome harboring linked mutations across the length of Gag-Pol, including one canonical drug resistance mutation that was not previously detected via clinical drug resistance testing. This data underscores the utility of the MrHAMER platform to identify the genetic correlates of intra-host viral evolution in response to antiviral therapy and immune pressure, with implications in the identification of interactions of novel targets that can be modulated for therapeutic benefit. Our approach is also widely applicable for sequencing of other RNA viruses and for adjacent fields that necessitate analysis of long-range genetic interactions including: RNA structure probing, haplotype phasing, organ/tissue transplantation compatibility (HLA typing), neutralizing antibody development, and gene expression of splice variants.

## Results

### Optimization and validation of RT-PCR conditions to minimize template switching and amplification bias while ensuring sufficient cDNA yield from low viral load samples

Given the limited viral load in clinical samples of patients undergoing antiviral therapy [28, 29], both RT and PCR conditions need to be optimized to obtain sufficient yield during reverse transcription and minimize template switching artifacts during PCR amplification. Conveniently, expected RNA input in clinical samples are several orders of magnitude lower than the levels at which RT template recombination becomes detectable via NGS [30]. We first focused on optimizing Reverse Transcription (RT) conditions by choosing a suitable priming strategy and testing for cDNA yield with limiting amounts of RNA inputs. A gene-specific primer (4609bp) was designed to target the highly-conserved region at Vif splicing junction in HIV RNA and used to generate a 4.6kb cDNA product that covers the Gag-Pol region and contains a synthetic priming site for downstream PCR amplification. This priming strategy was used to reverse-transcribe 100,000, 30,000 and 10,000 HIV RNA genome copies with either OneScript Plus RT (ABM) or SuperScript IV RT (ThermoFisher), followed by PCR amplification for yield evaluation. OneScript Plus, showed the greatest yield of full-length Gag-Pol cDNA compared to SuperScript IV at all tested RNA inputs **(Supplementary Figure 1)**. Following the establishment of suitable RT conditions, we proceeded to optimize PCR conditions with an emphasis on minimizing amplification artefacts.

Template switching is a known source of artefactual recombination when generating PCR-amplified cDNA for sequencing of HIV [31, 32] and other RNA viruses. To counteract this major PCR artefact we adapted established emulsion PCR (emPCR) protocols [33, 34] and systematically tested a range of conditions using a direct readout of template switching, where HIV RNAs containing 8-bp barcodes at either 5’ or 3’ ends are mixed at equal proportions, put through RT-PCR, then sequenced with ONT to count recombinant sequences containing both barcodes (or no barcodes) **(Supplementary Figure 2)**. The isolation of cDNA templates in their own emulsion droplets reduces the chances of artefactual recombination, with an added benefit of controlling for PCR sampling biases. Given that the amount of reactants in each emPCR droplet is restricted, and template switching has been previously shown to occur in latter amplification cycles [30, 35], two serial amplifications with reduced PCR cycles were adopted to minimize amplification bias and ensure preservation of long-range linkage. We determined emPCR dramatically reduced template switching rates compared to bulk PCR **(Figure 1A)**. Additional variables that reduced template switching include adoption of size selection between the emPCR reactions (that enriches for full-length product), and reducing the number of PCR cycles. The combined use of emPCR, reduced cycles, and size selection reduced template switching from a high of ∼42% in Bulk PCR to less than 1% (0.88±0.22%). Importantly, these optimized emPCR conditions were found to yield sufficient cDNA for downstream sequencing when starting with 5,000 copies of viral RNA **(Figure 1B)**.

**Figure 1.**
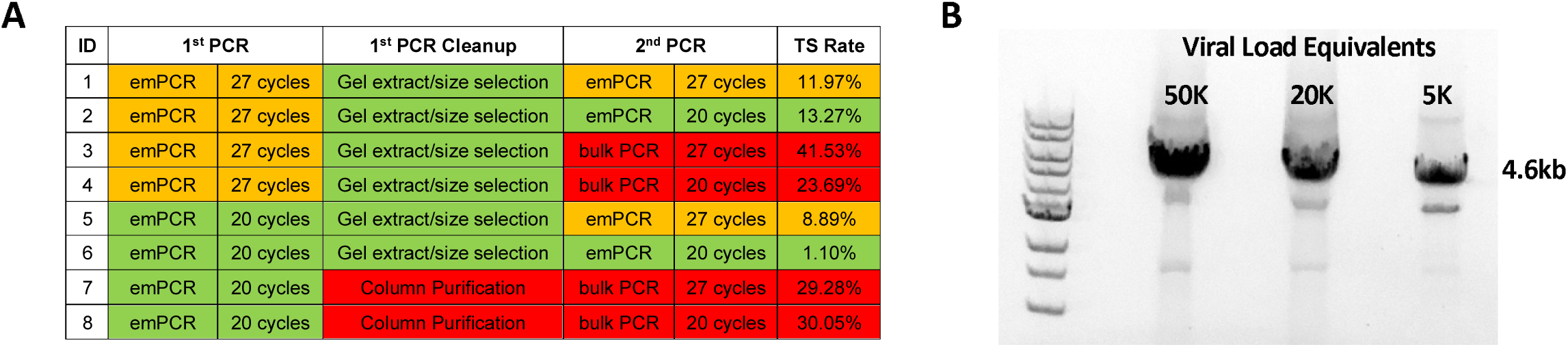
Optimization of RT-PCR conditions to preserve long-range linkage and ensure sufficient cDNA yield from low RNA inputs. (A) PCR amplification and sample handling variables tested, and their resulting template switch rates. (B) Agarose gel electrophoresis of amplified cDNA products using RNA input amounts corresponding to 50,000, 20,000, and 5,000 viral RNA copies, under optimized conditions.

### Nanopore sequencing of Gag-Pol cDNA to high accuracy using MrHAMER

Sequencing of viral cDNA to high accuracy using PacBio or Oxford Nanopore requires a number of adaptations given the 5-10% single-pass error rates in these single-molecule sequencing platforms [36]. A number of methods have been developed that increase the accuracy of these platforms, all sharing the general tactic of generating multiple repeats of each template and using these repetitive units for error correction during the computational processing step. Two such approaches were initially published, R2C2 [37] and INC-Seq [38], both resulting in 2-3% error rates in Nanopore platforms, but they are optimized for spliced transcript detection, or use chemistries that are no longer supported. Moreover, both these approaches do not have a way to curb or detect template switching events between similar templates, as would be present in the viral swarm of circulating HIV RNAs. A more recent Nanopore-based error-correction scheme uses computational filtering via dual barcodes to control for template switching, but the overall methodology has not been validated for viral genomics applications [39].

We therefore developed a library preparation and bioinformatic pipeline that generates concatenated sense and antisense repeats of a single molecule that are used by a downstream bioinformatic pipeline for error correction purposes. This error-correction workflow generates concatemers via a rolling circle amplification (RCA). Briefly, hairpin adapters are ligated to end-prepped cDNA using the NEBNext Ultra II kit **(Figure 2A)**. These hairpin adapters, analogous to those found in NEBNext and PacBio HiFi kits, contain an open region where a phosphorothioated primer can bind and initiate linear extension with EquiPhi29, generating concatemers with sequential repeats of sense and antisense strands **(Figure 2B)**. The ability to capture both strands of an originating single molecule is a differentiating feature in our approach, and can result in greater error-correction potential by providing the reverse complement of a problematic or error-prone *k-mer*. Moreover, the concatemers are generated in a dimethicone (DMF-A-6CS) based emulsion [40] which minimizes both sequence chimeras and non-specific Phi29 amplification, the latter being a persistent problem in isothermal amplification reactions [41]. Following second-strand generation and size-selection, the high-molecular weight (HMW) DNA is prepared for sequencing using standard Nanopore ligation kits for sequencing on R9.4.1 flow cells.

**Figure 2.**
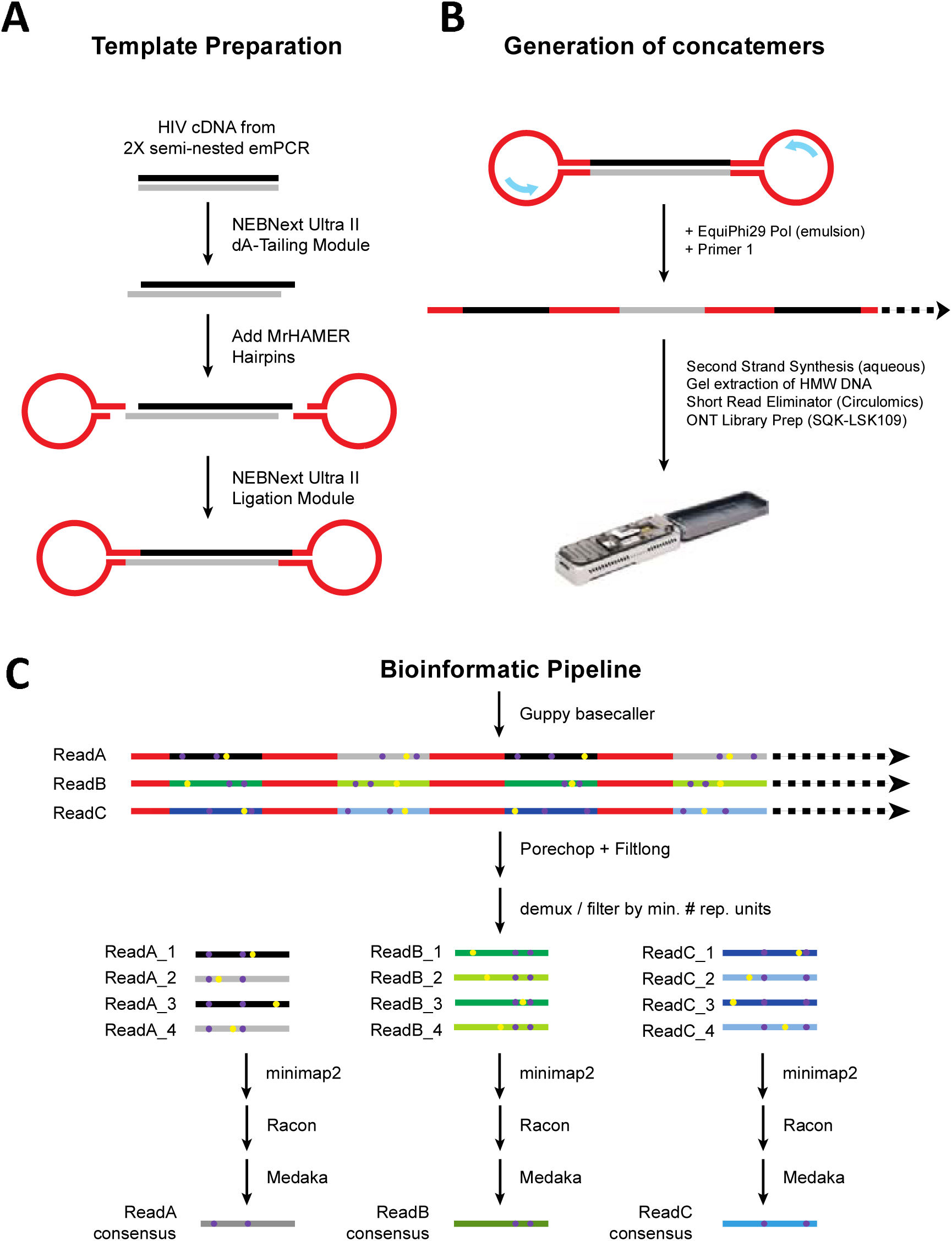
MrHAMER template preparation and bioinformatic pipeline. (A) Template preparation involves ligation of hairpin loop adapters to both sides of end-prepped amplified cDNA. (B) Concatemers are generated in emulsion using EquiPhi29 polymerase with a phosphorothioated gene specific primer, followed by second strand synthesis and size selection prior to ONT sequencing. (C) Bioinformatic pipeline involves breaking up concatemers into its constitutive sense and antisense repeats, followed by demultiplexing into single FASTQ files composed of repetitive units originating from a single concatemeric read. Each FASTQ file, is reference mapped to the HIV reference, followed by Racon and Medaka polishing, resulting in error-corrected FASTA files

The MrHAMER bioinformatic pipeline **(Figure 2C)**, leverages a combination of publicly available tools with custom python scripts for ease of use and modularity. Briefly, Porechop (https://github.com/rrwick/Porechop) is used to split concatemers based on the presence of MrHAMER hairpin sequence, the resulting reads file is then demultiplexed with a custom python script into FASTQ files for each originating single molecule and containing all concatemeric repeats, while filtering for a minimum number of repeating units. The next steps are parallelized and involve alignment of each FASTQ file to an HIV reference using minimap2, followed by polishing using the Racon (github.com/lbcb-sci/racon), and Medaka (github.com/nanoporetech/medaka) sequence correction packages. This results in high accuracy viral sequences in FASTA format, each constituting a polished single genome containing a full-length Gag-Pol sequence.

Reconstructed genomes show optimal coverage across the length of the Gag-Pol region **(Figure 3A)** without evidence of positional biases. Similar to previous RCA-type error correction methodologies, reductions for each error type (i.e. mismatch, deletion, insertion) is directly proportional to the number of repetitive units generated per single molecule **(Figure 3B)**. Overall, the MrHAMER pipeline reduces Nanopore error rate from ∼11% to 0.133% when 10 repetitive units are included in a single long read. This constitutes an 84-fold accuracy improvement over the baseline single-repeat error rate and yields median Phred quality scores of 28.76 **(Figure 3C)**. Additional error correction can be achieved by requiring a higher number of repetitive units, at the expense of overall number of single molecule genomes obtained per sample. Our computational pipeline is designed so this filtering criteria can be adjusted if greater number of genomes is necessary (at the expense of slightly lower accuracy). As expected, deletion rates remain the predominant contributor to the error profile of MrHAMER, particularly proximal to homopolymeric regions, which is consistent with the known characteristics of the R9.4.1 Nanopore chemistry **(Figure 3D) [39]**. In addition to using IVT RNA inputs, MrHAMER was validated for the ability to obtain accurate genomes from live NL4-3 virus, with error rates being reduced from 9.628% to 0.111% and with quality scores of 29.55 **(Supplementary Figure 4)**, which approaches quality values observed in Illumina Sequencers.

**Figure 3.**
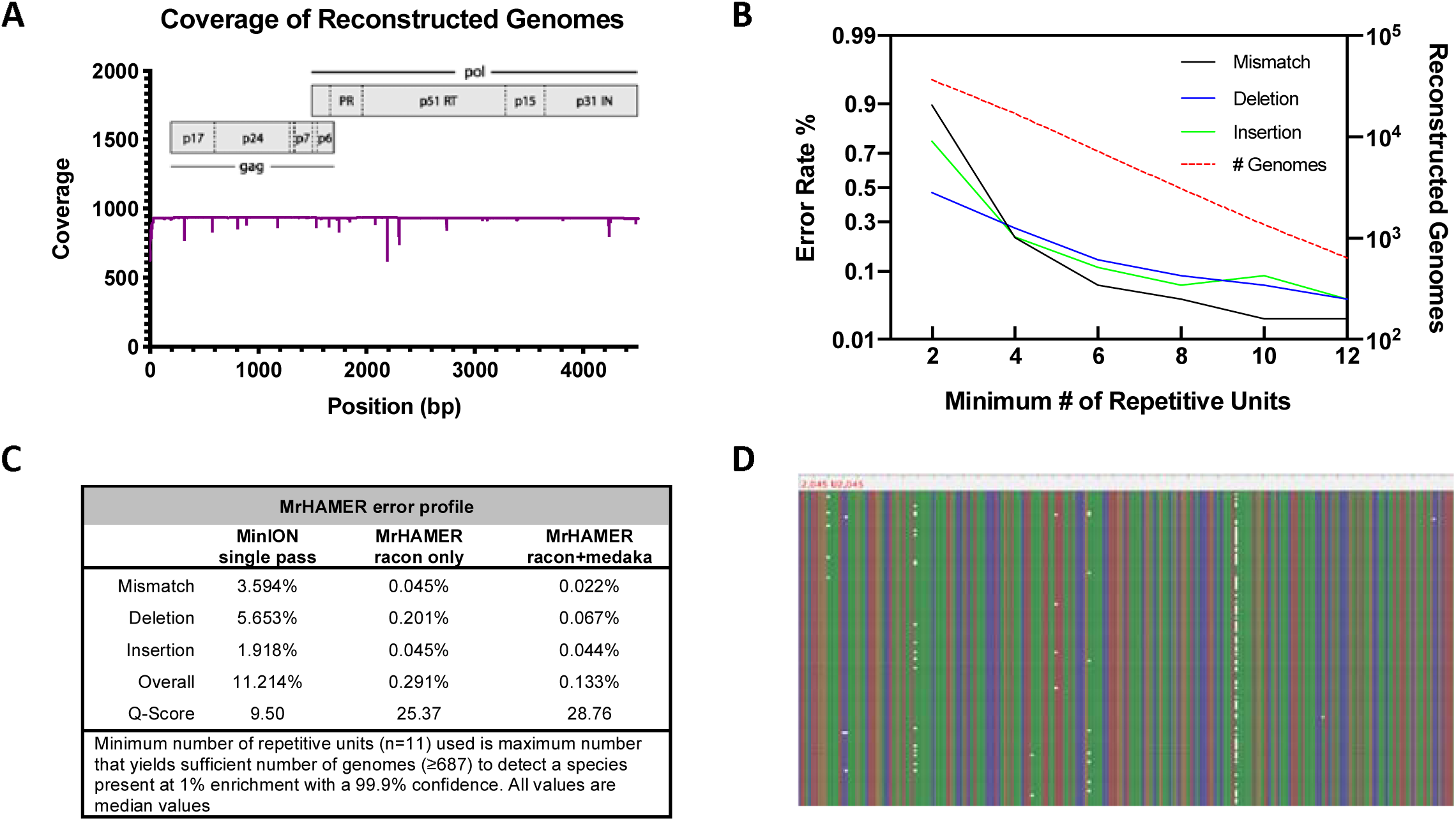
MrHAMER generates highly accurate single-genome reconstructions with even coverage along Gag-Pol region. (A) Coverage of reconstructed genomes aligned to HIV reference shows even coverage along length of Gag-Pol region (B) Extent of error-correction positively correlates to the number of repetitive units used and negatively correlates to MrHAMER throughput as measured by number of reconstructed genomes (i.e. number of error-corrected Gag-Pol reads). A minimum number of eight to ten repetitive units is a reasonable compromise between error rate and throughput. (C) MrHAMER reduces the error rate of ONT sequencing from ∼11% on a single pass, to 0.133% when using an optimal number of repetitive units for correction, resulting in quality scores approaching parity with Illumina sequencing. (D) Sequence alignment of reconstructed genomes was visualized with Tablet and shows deletions constitute the bulk of errors that remain after error correction, and they are invariably located proximal to homopolymeric or low-complexity regions.

### MrHAMER preserves the original proportion of mutation pairs in an *ad hoc* complex mixture and allows detection of linked mutations present at ≤1% frequency

To validate whether MrHAMER can accurately detect the positions and nucleotide identity of linked mutations at their expected frequency, we generated two proviral constructs derived from a pSG3.1 HIV strain and each containing mutation pairs. *In vitro* transcribed RNA was generated from these constructs and mixed in a 75:24 proportion, with IVT RNA from unmodified ‘wild-type’ pSG3.1 construct mixed in at 1% frequency to generate an artificial viral population mix. Input RNA mixture was reverse-transcribed and PCR-amplified following the MrHAMER pipeline described above and requiring at least 10 repetitive units per genome. To reliably determine high confidence linked mutations, MrHAMER outputs are analyzed with the Co-Variation Mapper (CoVaMa) pipeline [17]. MrHAMER sequencing preserved the original proportion of linked mutations, with only a slight overrepresentation of the ‘wild type’ background originally mixed in at 1% frequency **(Figure 4A)**. More importantly, the error-correction of MrHAMER was suitable for detection of all linked mutations at their expected position **(Figure 4B)** and with proper nucleotide identity **(Supplementary Table 4)**. This data provides critical validation that MrHAMER is able to accurately identify linked mutations present in a complex mixture with sufficient sensitivity and signal to noise ratio.

**Figure 4.**
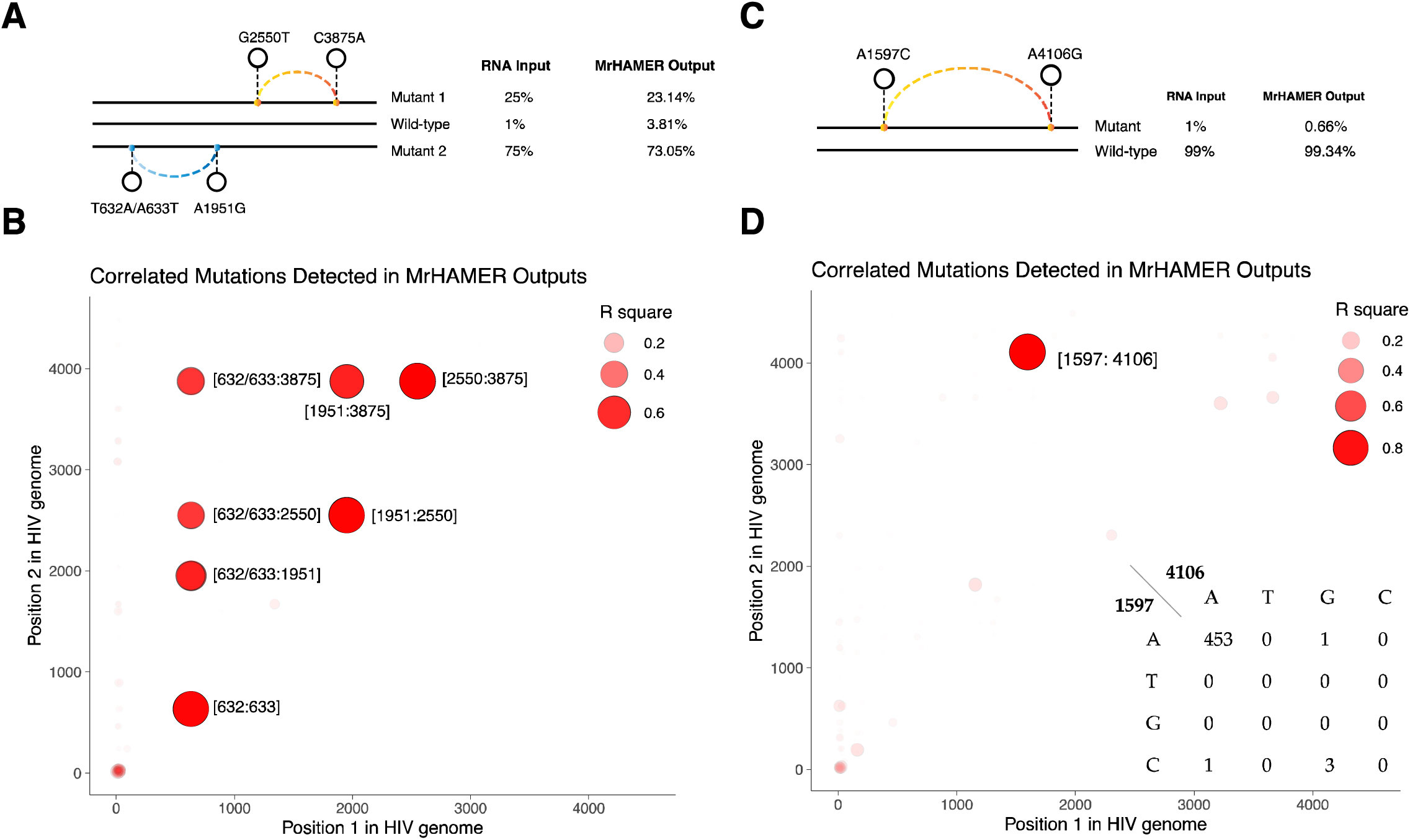
CoVaMa accurately detects linked mutations in complex mixtures from MrHAMER outputs, with ability to detect linked mutations present at ≤1% enrichment. (A) Two mutants containing distant linked SNVs, and one ‘wild-type’ RNA were admixed at the indicated proportions to generate a synthetic mixture of strains each containing specific linked mutations. MrHAMER preserved the proportion of this complex mixture as analyzed via variant-calling analysis. (B) MrHAMER outputs were analyzed with CoVaMa, which successfully detected all co-variant pairs with all expected coordinates showing high R^2^ values. To determine sensitivity, we mixed single mutant containing linked SNVs at 1% enrichment with ‘wild-type’ HIV RNA present at 99% enrichment. (C) MrHAMER preserved the proportion of the mixture and (D) successfully detected this rare variant with an R2 value of 0.8.

To test the ability of MrHAMER to capture low-frequency linked-mutations present at ≤1%, an additional proviral mutant was designed to generate IVT RNA, then mixed at 1:99 ratio with ‘wild-type’ IVT RNA. After MrHAMER sequencing, linked mutations are detected at 0.66% frequency, which is sensibly close to the admixed value of 1% **(Figure 4C)**. CoVaMa shows the mutation pair is detectable with a high R^2^ value of ∼0.8 **(Figure 4D)**, and with contingency tables showing sensitivity of ≥99.6% **(Figure 4D, inset)**. These data provide full validation of the ability of MrHAMER to yield an accurate readout of linked mutations in complex mixtures of viral species, with enough sensitivity to detect linked mutations present at <1% enrichment within the viral pool.

### Sequencing of longitudinal HIV isolates from a patient undergoing virological failure shows clonal expansion of a pre-existing Gag-Pol genome harboring linked drug-resistant mutations

Having validated the MrHAMER sequencing pipeline using *in vitro* control datasets, we proceeded to sequence clinical samples to characterize genetically linked variants in the HIV Gag-Pol region that correlate with cART failure. For this proof of concept, we used sequential plasma samples from a single patient collected before and after antiretroviral therapy, with the latter sample collected during viral load rebound (i.e. virological failure). The operating assumption was that linked mutations enriched during virological failure provide increased viral fitness in the presence of antiretrovirals. Viral sequences obtained prior to cART initiation can then be interrogated for the presence of these linked mutations to delineate possible mechanisms of viral evolution that resulted in eventual antiretroviral failure.

Blood plasma was obtained from a single patient at two time points: prior to initiation of cART therapy (i.e. Treatment Naïve), and following multiple instances of therapy failure to cART (i.e. Virological Failure, VF) **(Supplementary Table 1)**. The patient’s initial treatment course included a protease inhibitor coupled to nucleoside (NRTI) and non-nucleoside (NNRTI) reverse transcriptase inhibitors. Genotypic drug resistance testing prior to cART initiation showed no major DRMs were detectable to any of the used drugs classes **(Supplementary Table 2)**. Antiretroviral therapy regimen was subsequently changed, each time to a cocktail of three NRTIs **(Figure 5A)**. Both therapy changes failed to prevent subsequent viral rebounds, with samples from the second NRTI-related viral rebound used for sequencing as a virological failure (VF) timepoint. MrHAMER sequencing of these isolates **(Supplementary Table 3)** yielded 1002 and 1270 Gag-Pol sequences for Naive and VF samples respectively (when using ≥10 repetitive units). Given the number of reconstructed genomes obtained per timepoint and the high intra-host diversity, we used CliqueSNV to extract statistically linked mutations and cluster them into full-length Gag-Pol haplotypes that shared patterns of SNVs [42]. This cuts through the biological noise generated by the inherently high mutation rate of the virus and allows identification of clusters (and their associated abundance) containing shared SNV patterns throughout the length of Gag-Pol. Using CliqueSNV, forty one Gag-Pol haplotypes were detected in the Naïve sample with the two most abundant haplotypes not exceeding 5% of the population. Conversely, twelve haplotypes were detected in the VF sample with the two abundant haplotypes having a combined frequency of 52%. The significant ∼3-fold reduction in the number of haplotypes following virological failure, along with the 10-fold enrichment of haplotype abundance is consistent with a reduction in Shannon Diversity Index from 3.51 to 2.02, and suggests the presence of a drug-pressure induced selective sweep that results in a predominant viral strain [26]. Since CliqueSNV generates a consensus sequence for each haplotype cluster, Naïve and VF haplotype FASTA sequences were concatenated, and a multiple sequence alignment (MSA) was performed, followed by phylogenetic analysis **(Figure 5B)**. Consistent with a strong selective sweep, phylogenetic analysis revealed that all the viral strains in the VF samples likely arose from a particular set of ancestor haplotypes in the drug-naive viral population (naïve haplotype IDs 24 and 56). Moreover, the visibly long branch length in the VF clade suggests strong genetic divergence due to selective pressure from cART treatment.

**Figure 5.**
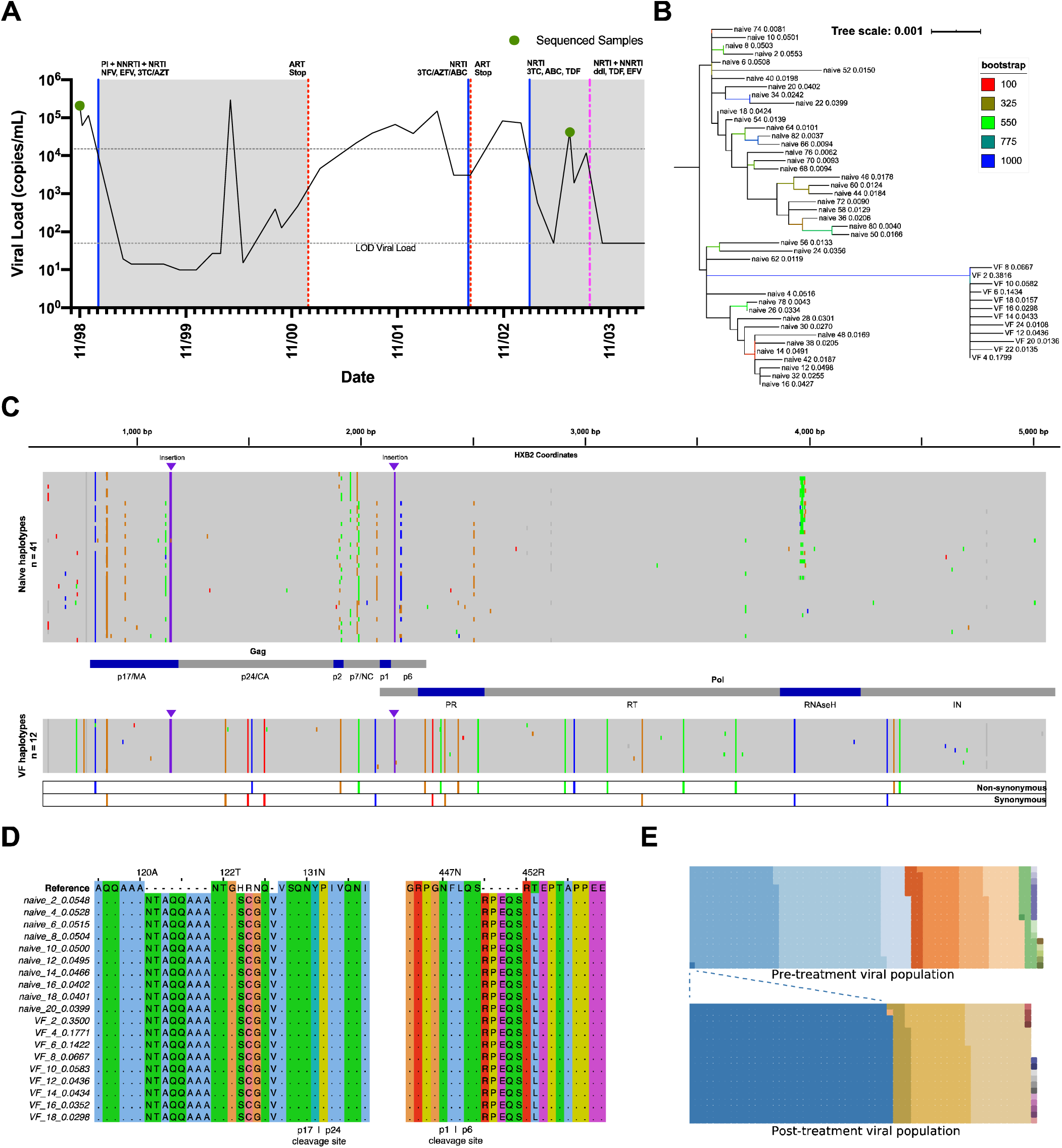
MrHAMER identifies genetically linked mutations in the HIV Gag-Pol region and reveals the clonal expansion of a pre-existing Gag-Pol genome harboring linked drug-resistant mutations in patient undergoing virological failure. **(A)** Viral load and Treatment history of patient undergoing virological failure. Naïve and virological failure (VF) samples are collected as shown in green dots. Antiretroviral therapy periods are shaded in grey **(B)** 41 and 12 haplotypes are respectively obtained from clustering of single-molecule Naïve and VF Gag-Pol reads based on shared mutation patterns. A phylogenetic tree is built from the MSA of these Gag-Pol haplotypes sequences using the Neighbor joining algorithm and 1000 bootstrap replicates, with the branch length proportional to genetic distance. The phylogenetic tree is graphed with iTOL and rooted to the Naïve consensus sequence. Each haplotype branch shows: sample type, haplotype ID, and haplotype frequency within each Naïve or VF sample. **(C)** Haplotype sequences obtained from Naïve and VF samples are mapped to the Naïve consensus sequence and visualized with IGV. 25 mutations significantly enriched during the virological figure within Gag-Pol ORF, with 60% resulting in non-synonymous mutations (including canonical M184I in RT region) and 40% resulting in synonymous mutations. Two insertions in the Gag region, which result in duplication of AA motifs are highlighted by purple arrowheads. **(D)** MSA outputs of 10 most enriched haplotypes in each Naïve and VF sample shows insertions present upstream of p17 | p24 and downstream of p1 | p6 cleavage sites, and **(E)** Highly-accurate single-molecule Gag-Pol sequences from Naïve and VF samples are divided into subgroups based on their sequence identity at 25 positions that where enriched during VF. A waffle plot is generated for each sample, with each square constituting a single molecule sequence with unique nucleotide identities at these 25 positions. Color across sample type are color-matched for identification of clonal

For the next steps, analysis of mutation patterns in Gag-Pol genomes across longitudinal samples is performed with respect to consensus viral sequence present prior to treatment initiation. SNVs in the Gag-region (particularly in p17, p2, p7, and p6) and in Protease and RNaseH Pol regions contributed the most to population diversity in the drug-naive sample **(Figure 5C)**. Comparison between two longitudinal samples showed VF-associated enrichment of a viral strain containing 25 linked SNVs within the Gag-Pol open reading frame, with at least one SNV resulting in the major M184I surveillance drug-resistance mutation in RT domain **(Table 1)**. The enriched SNVs associated with therapy failure result in both synonymous and non-synonymous mutations, with the latter constituting 60% of SNVs and being mostly found within the RT/RNAseH domain (as consistent with single-class NRTI cocktail used prior to VF timepoint). In addition, alignment of full-length Gag-Pol Naïve and VF haplotypes revealed two insertions in Gag that existed in both Naïve and VF samples, which suggest a transmitted founder strain contained these features which are likely energetically neutral and are retained after drug exposure. The first insertion was located directly upstream of the p17/p24 cleavage site and results in a duplication of a QQAAANT motif, while second insertion is located downstream of the p1/p6 cleavage site and results in a duplication of a QSRxE motif **(Figure 5D)**.

**Table 1.** Single nucleotide variants present in most enriched VF haplotype.

| Nucleotide<sup>1</sup> | Type | Mutation<sup>2</sup> | Protein | Domain |
| --- | --- | --- | --- | --- |
| G816C | Non-synonymous | R9S | Gag | p17 |
| A867G | Synonymous | - | Gag | p17 |
| A1395G | Synonymous | - | Gag | p24 |
| A1497T | Synonymous | - | Gag | p24 |
| A1514C | Non-synonymous | N242T | Gag | p24 |
| C1572T | Synonymous | - | Gag | p24 |
| A1909G | Non-synonymous | I374A | Gag | p2 |
| C1990A | Non-synonymous | L401I | Gag | p7 |
| T2064C | Synonymous | - | Gag | p7 |
| A2286G | Non-synonymous | T12A | Protease | - |
| C2319T | Synonymous | - | Protease | - |
| C2357A | Non-synonymous | D35E | Protease | - |
| A2375G | Synonymous | - | Protease | - |
| A2436G | Non-synonymous | I62V | Protease | - |
| G2522A | Synonymous | - | Protease | - |
| G2913A | Non-synonymous | E122K | RT | - |
| T2953C | Non-synonymous | I135T | RT | - |
| G3101A | Non-synonymous | M184I | RT | - |
| T3257G | Synonymous | - | RT | - |
| C3439A | Non-synonymous | A297E | RT | - |
| G3672A | Non-synonymous | V375I | RT | - |
| A3935C | Synonymous | - | RNAseH | - |
| T4349C | Synonymous | - | IN | - |
| A4379G | Non-synonymous | I50M | IN | - |
| G4405A | Non-synonymous | G59E | IN | - |
| <sup>1</sup> Nucleotide positions based on HXB2 coordinates |  |  |  |  |
| <sup>2</sup> Amino acid mutation coordinates for Gag are for polyprotein form while coordinates for all Pol proteins are for each cleaved product |  |  |  |  |

To have a more granular understanding of the potential mechanisms that gave rise to the selective sweep that enriched for 25 specific SNVs during virological failure, we interrogated the specific nucleotide identities at each of these 25 genome positions for each of the Gag-Pol sequences obtained in Naïve and VF samples. The positions of these 25 enriched mutations were selected, and the viral population in each sample was divided into several subgroups identified by their unique combination of nucleotides at these positions. Consistent with the CliqueSNV outputs, the Naive sample showed higher cluster diversity while the VF sample was more homogenous **(Figure 5E)**. Strikingly, a clonal expansion of a single viral genome was identified, where this rare species is enriched from ∼0.1% at the treatment Naïve stage to 53.4% at virological failure and contains all 25 SNVs that are predominantly enriched in VF samples, including the non-synonymous mutation that results in the major M184I DRM. This shows our assay was sensitive enough to capture a rare pre-existing Gag-Pol sequence harboring a major DRM (and several other linked SNVs) that is enriched during virological failure, but was previously undetectable via clinical genotypic drug resistance testing prior to ART initiation. This suggests that the selective sweep that resulted in VF in this patient was not necessarily caused by recombination of pre-existing (or *de-novo*) SNVs, but was caused by a pre-existing virus containing all linked mutations and DRMs that was selected due to a pleiotropic or functional epistatic effect.

## Discussion

In this study, we introduce an end-to-end sequencing pipeline that outputs highly-accurate HIV genomes covering the entire Gag-Pol region from clinical samples, and captures long-range co-evolved mutations while providing full readout of genetic background of each virus. Our MrHAMER approach is facilitated by validated RT-PCR conditions to obtain accurate and unbiased amplified cDNA from samples with minimal viral load, and by a template preparation and bioinformatic pipeline that allows nanopore sequencing of this cDNA to high accuracy. As a proof of concept, we apply the MrHAMER pipeline to sequence longitudinal HIV isolates from a patient undergoing therapy failure and identify a hard selective sweep from a pre-existing drug-resistant Gag-Pol genome that correlates with VF.

The rapid and continued developments in the Nanopore computational processing space have greatly facilitated the development of our MrHAMER approach [36]. Recent advances in recurrent neural network-based basecalling and polishing have enabled iterative improvements in the error-correction potential of our pipeline by increasing both accuracy and throughput. Namely, our transition from version 3.4.5 to 3.6.0 of the Guppy basecaller, and the use of updated models in Medaka polisher, have resulted in 2-fold decreases in the MrHAMER error-rate, a 50% increase in the number of reconstructed genomes for a given sequencing run, and higher signal-to-noise ratio in CoVaMa outputs. In the near term, the development of run-length encoded and convolutional neural network [43, 44] basecallers, show potential in increasing single-pass accuracy proximal to homopolymeric regions, resulting in further accuracy improvements of our single molecule viral genomes. Further accuracy increases in the longer term are likely to be facilitated by direct analysis of the raw signal-level data of the sense and antisense repeats produced by MrHAMER, either by leveraging pair consensus or multidimensional basecalling [45], or signal-level consensus polishing via Nanopolish [46]. Any of these accuracy improvements can be implemented in MrHAMER without having to re-sequence samples, allowing reanalysis of raw data to even greater accuracy and throughput.

Our MrHAMER approach currently provides highly accurate single-molecule viral RNA sequences of ∼5kb in length, with longer sequences requiring a tiled amplicon approach. There are three constraints that dictate our current limit in the length of reconstructed genomes: (1) RT processivity limitations for RNA templates >5kb, (2) length of concatemers produced via Phi29, and (3) number of repetitive units required for sufficient error correction given current basecalling technologies. Reverse transcription of RNA templates beyond 5kb remains a methodological challenge with most commercially-available RT enzymes. This constraint is driven by processivity limitations of these MMLV-/AMV-derived RTs [47] and is exacerbated by highly structured regions in viral RNAs, particularly in HIV. Alternatively, novel RTs with increased processivity [48, 49] might result in sufficient cDNA from longer RNA inputs; however, these enzymes need to be validated for their performance when using the limiting RNA inputs that are typically found in clinical isolates. With regards to the production of concatemers containing sufficient repeats for error-correction, this is predominantly a variable of the processivity of the Phi29 variants used for RCA within emulsion. For example, obtaining sufficient error correction from full-length 9.2 kb HIV cDNA would push the Phi29 polymerase to generate a concatemer of double the length of what would be required for our current 4.6 kb Gag-Pol genomes. Since our MrHAMER approach already uses a highly-processive mutant of Phi29 (EquiPhi29, ThermoFisher), likely opportunities for optimization hinge on maximizing emulsion mono-dispersity coupled to an increase in the overall size of the emulsion droplets to ensure homogeneous and optimal availability of reagents. Conversely, continued improvements in basecalling and polishing platforms might obviate the need to increase length of concatemers when targeting RNA >5kb by allowing use of less repetitive units for comparable error-correction.

An immediate application of our sequencing approach is to interrogate the viral evolution mechanisms of drug resistance in patients going through virological failure, with particular emphasis on understanding pleiotropic or functional epistatic pathways that precipitate this clinical outcome. Pleiotropic and functional epistatic effects have been explored in the past within the context of HIV drug resistance; however, the methodologies did not allow the direct observation of co-evolved mutation pairs beyond the short read-lengths afforded by Illumina sequencing, or relied on imputing and inferring co-evolution via bioinformatic or computational approaches which were shown to be unreliable given the high intra-host genetic diversity in HIV [18]. With the ability of MrHAMER to capture distant co-evolved mutations while providing a full readout of the genetic background of each virus, we set out to establish possible pathways and their associated genetic correlates that drive virological failure in a patient. The resulting proof of concept study using longitudinal HIV isolates revealed the clonal expansion of a pre-existing Gag-Pol genome harboring linked SNVs associated to the M184I NRTI drug resistance mutation (DRM). Drug resistance mutations, including M184I NRTI mutations, have been shown to negatively affect replicative capacity of the virus, thus the viral strain harboring these DRMs would tend to revert to wild type or acquire compensatory mutations [50, 51]. The acquisition of compensatory mutations could prevent reversion, either by restoring protein function in the presence of inhibitor, or stabilizing the structure their enzymatic substrates and contribute to the establishment and maintenance of viral fitness [52]. Two lines of evidence suggest the linked mutations associated with M184I play a compensatory role in restoring viral replicative capacity of this mutant viral strain: (1) the vast majority (∼90%) of linked mutations were not individually present (beyond a single molecule) at the treatment naïve stage and thus not favored in the absence of drug pressure, and (2) linked mutations were enriched in response to drug pressure instead of reverting to WT. Our methodology underscores that the genetic background of a viral strain harboring DRMs could play an essential role in shaping treatment outcomes and provides a starting point to study long-range epistasis in the HIV and other viruses. Our observation regarding co-evolved mutation sets that correlate with drug resistance needs to be thoroughly validated, with particular care in testing the effect of full Gag-Pol viral blocks/strains (not just individual SNVs) in relevant biochemical and *in vitro* assays.

In addition to SNVs that resulted in amino acid mutations, MrHAMER also revealed a network of linked synonymous mutations that correlate with virological failure and constitute up to 44% of all enriched SNVs. These synonymous mutations are evenly spread throughout Gag-Pol and are not pre-existing (beyond single precursor genome), with their enrichment after drug pressure pointing to presumed fitness advantage. This finding is quite surprising given the prevailing consensus that synonymous mutations may disrupt functionally important RNA structures or cis-regulatory elements that overlap with open reading frames (ORFs) [53, 54]. Recent global synonymous mutagenesis studies confirmed that synonymous mutations do have a deleterious effect on replication competence, infectivity, and splice site usage, but these effects are constrained by proximity of these SNVs to splice donor/acceptor sites[55] where dynamic RNA structure modulates accessibility of these sites to spliceosomal components[56]. In our samples, three of the eleven synonymous SNVs are located within the N-terminus of Gag, a ‘block’ previously shown to be critical for viral fitness in synonymous mutagenesis studies. The fact these three mutations are enriched without reverting to WT in response to drug pressure, suggests these synonymous variants result in increased fitness or are positively selected due their effect in replicative capacity. The remaining eight synonymous mutations are outside of these previously interrogated sites of spliceosomal regulation, which suggest they are energetically neutral or, alternatively, might be part of yet uncharacterized cis-regulatory elements or ribonucleoprotein interfaces. The exact contribution of these synonymous mutations needs to be investigated to address whether they play a compensatory role to other non-synonymous mutations, or if they are providing benefits in viral fitness that are independent from amino acid changes.

In summary, we developed a nanopore-based long-read viral sequencing methodology that yields highly-accurate single-molecule sequences of circulating virion RNA from clinical samples. We validate and demonstrate the utility of our approach by sequencing HIV RNA covering the entire Gag-Pol region from longitudinal samples from a patient undergoing virological failure. We believe the MrHAMER pipeline will alleviate the major bottleneck in the direct observation of long-range co-evolution in HIV and other RNA viruses at the single-molecule level. The granular view of intra-host diversity provided by our pipeline will facilitate scalable investigation of genetic correlates of resistance to both antiviral therapy and immune pressure, and enable the identification of novel host-viral and viral-viral interfaces that can be modulated for therapeutic benefit.

## Acknowledgements

The authors wish to thank John Coffin (Tufts), Stephen Hughes (NCI), and Mary Kearney (NCI) for helpful comments and advise in early stages of the project. Special thanks to Steven Head and Phil Ordoukhanian from the TSRI NGS Core for helpful insights and reagents. Thanks Brian Paegel (UCI) for providing critical reagents and expertise on emulsion chemistry. Thanks Ali Torkamani (TSRI) for discussions on bioinformatic methodologies. Thanks to Stosh Ozog (TSRI) for providing NL4-3 viral supernatants and Michael Zwik (TSRI) for providing pSG3.1 proviral vector.

## Author Contributions

CMG, ALR, and BET conceived and conceptualized the study. CMG developed assay portion of study. CMG, DJM, and ALR conceived the bioinformatics pipeline. CMG, SW, ALR, and BET wrote the manuscript. CMG and SW sequenced and processed samples. CMG and ALR optimized RT and PCR workflows. CMG and SW developed control datasets. CMG and SW validated methodology. CMG, SW, DJM, and ALR developed bioinformatic tools. CMG, SW, ALR, and BET analyzed and interpreted data. CMG and SW generated figures. SJL and DMS selected and provided patient samples. All authors discussed content and revised final manuscript draft.

## Data availability

Raw sequencing data will be available on Sequence Read Archive (SRA) upon acceptance of publication.

## Code availability

Python scripts will be available on Github upon acceptance of publication.

## Methods

### Generation of plasmids for In Vitro Transcription of HIV RNA

To generate a plasmid for in vitro transcription of HIV RNA, the HIV insert from the pSG3.1 strain [57] was PCR amplified with Q5 HotStart Master Mix in two fragments, with an overlap in the PR locus to add a D25A mutation and both an EcoRI/T7 promoter and PolyA/BamHI sites added at the 5’ and 3’ ends of insert. A pUC19 backbone was PCR amplified with overlaps to the T7 promoter site at the 5’ of insert and PolyA tail at the 3’ end. PCR-amplified insert and vector fragments were assembled with NEBuilder HiFi DNA Assembly kit (E2621S) and plated on an LB-Amp plate. Single colonies were grown, mini prepped, and sequenced to verify plasmid identity and orientation of all fragments. For nomenclature purposes, this sequence is referred to as a ‘wild-type’ strain throughout.

Additional mutants were designed based on this SG3.1 ‘wild-type’ background, all containing the following mutation pairs: T632A/A633G with A1951G, G2550T with C3875A, A1597C with A4106G. These mutations were added via PCR amplification with primers that generate overlaps for subsequent NEBuilder HiFi assembly. All modified plasmids containing mutation pairs were grown from single colonies and sequenced to verify sequence identity.

### In Vitro Transcription of HIV RNA

HIV plasmid is treated with T5 exonuclease (NEB M0363S) to digest any fragmented vector, and DNA cleaned with Monarch PCR & DNA Cleanup Kit (NEB T1030S). Resulting supercoiled plasmid is linearized at the 3’ end of the PolyA tail using BamHI-HF (NEB R3136S), and checked for reaction completion by running on agarose gel. Linearized plasmid is DNA cleaned, and eluted in nuclease free water. Standard RNA Synthesis was carried out with the HiScribe T7 High Yield RNA Synthesis kit (NEB E2040S) for 1.5 hours according to the manufacturer’s instructions, using 500ng-1000ng of linearized plasmid as input, followed by DNase I digestion as instructed. RNA is purified using RNA Clean & Concentrator -5 kit (Zymo Research R1013) and eluted in nuclease free water. RNA samples were serially diluted in order to arrive at the desired number of input RNA molecules. When using a complex mixture of samples, RNA species were first mixed at high concentration at the right proportion, followed by serial dilutions to the appropriate RNA molecule number.

### Primer and Oligo Sequences

#### Reverse Transcription

RT_4609bp_Gal10pro-F:

GGTGGTAATGCCATGTAATATGNRNYNRNYNRNYNRNNNNNAGATCGGAAGAGCGTCGTGTCAC CTGCCATCTGTTTTCCATA

#### PCR amplification

1.U5.B1F: CCTTGAGTGCTTCAAGTAGTGTGTGCCCGTCTGT

2.U5.B4F: AGTAGTGTGTGCCCGTCTGTTGTGTGACTC (Nested within 1.U5.B1F)

Gal10-proF: GGTGGTAATGCCATGTAATATG

#### MrHAMER template prep and concatemer generation

MrH_Hairpin: 5’-Phos-TCTCTCTCTTTTCCTCCTCCTCCGTTGTTGTTGTTGAGAGAGAT

MrH_primer: AACGGAGGAGGAG*G*A (* denotes phosphorothioated position)

### Reverse Transcription

Reverse transcription is carried out with OneScript Plus Reverse Transcriptase (ABM, G237) in a 20µL volume with the following components and final concentrations: 1X Reaction Buffer, dNTPs (0.5 mM), RNAseOUT (2U/µL), RT_4609bp_Gal10pro-F gene specific primer (0.2 µM), RNA input (<5 µg), and OneScript Plus RT (10 U/µL). Primers are initially annealed to template RNA in the presence of dNTPs, by heating to 65°C for 5 min, followed by snap cooling to 4°C for 2 mins. After snap cooling, the rest of the components are added, followed by reverse transcription for 1.5 hours at 50°C. Reactions are stopped by heat inactivation at 85°C for 5 mins. Any residual RNA is digested by adding 1 µL each of RNaseI_f_ (M0233S) and RNAseH (M0297S). Reaction is cleaned with Monarch DNA Clean kit and eluted in 10 µL EB.

### Emulsion PCR

Aqueous phase is prepared in a 1.5 mL DNA LoBind tube to a final volume of 50 µL with the following components and final concentrations: cDNA input, 1X Phusion GC Buffer, 0.2 mM dNTPs, 0.5 µM of each primer, 0.5 mg/mL BSA, and 0.02U/µL Phusion U Hot Start DNA Polymerase (F555S). Oil/Surfactant is prepared: 2% (v/v) ABIL em90 (Evonik Degussa GmbH) and 0.05% (v/v) Triton X-100 in Mineral oil[33]. 300 µL of Oil/Surfactant is added on top of the aqueous phase, then vortexed for 5 mins at max speed using Vortex-Genie 2. Emulsified components are aliquoted into PCR strip (each tube containing no more than 50 µL). Samples are thermally cycled through following program: 98°C (2 min), 98°C(10 sec)/62°C(30 sec)/72°C(7:30 mins) for 20 or 27 cycles, 72°C(5 min) and 4°C Hold. Following thermal cycling, emulsion is consolidated in a 2 mL DNA LoBind tube, and emulsion is broken by adding 700 µL Ethyl Acetate and vortexing for 5-10 seconds. 1 mL of DNA binding buffer (Monarch DNA Clean) is added followed by vortexing for 10 seconds. Tube is spun at 20K RCF for 2 mins, resulting in three phases. Aqueous phase, which settled in bottom of tube, is carefully aspirated, and transferred directly to the DNA clean column with cleanup washes proceeding as indicated on manufacturer’s protocol, followed by elution in EB buffer. Cleaned up DNA is ready for agarose gel electrophoresis with E-Gel EX 1% gel system. The lane ‘band’ containing fragments in between 3kb and 10kb size is excised and size selected via gel extraction with Monarch kit and elution in EB. Since two serial emulsions PCR are required, first emPCR uses 1.U5.B1F and Gal10pro-F primers, whereas second emPCR uses 2.U5.B4F and Gal1pro-F primers, with forward primers being nested for higher specificity [58]. Size selection with gel extraction is performed after each emPCR for increased specificity and preservation of long-range linkage.

### MrHAMER template preparation

Purified Double-stranded cDNA from 2X emPCR, is A-tailed and end-repaired using NEBNext Ultra II End Prep kit (E7546S) according to manufacturer’s instructions. The MrHAMER hairpin oligo at 15 µM concentration (in 10mM Tris/10mM NaCl) is self-annealed by incubating at 90°C for 2 min followed by snap cooling in icy water. MrHAMER hairpin is added to the NEBNext Ultra II Ligation Master Mix, which is added directly to end-prep reaction tube. Ligation is performed at 20°C for 15 minutes, followed by AMPure XP bead clean up at 0.45X volume of beads, and eluted in 0.1X TE. Unligated, or partially ligated hairpin adapters are digested with exoIII and exo VIII in CutSmart Buffer for 30 mins, followed by two serial AMPureXP cleanups at 0.45X bead volume.

### Generation of concatemers from MrHAMER templates

Concatemers are first generated from MrHAMER templates via EquiPhi29 (ThermoFisher, A39390) amplification in emulsion. The emulsifier consists of 74.4% (v/v) DMF-A-6CS, 21.1% (v/v) Mineral Oil, and 4.5% (v/v) KF6038 and is made fresh for each reaction and placed on ice. Aqueous phase of EquiPhi29 reaction is assembled in a volume of 50 µL with following components and final concentrations: MrHAMER template (at least 3 ng), 1X EquiPhi29 Buffer, phosphorothioated MrHAMER_PS primer (5µM), dNTPs (1mM), DTT (1mM), Yeast Inorganic Phosphatase (0.025 U), BSA (0.5 mg/mL), EquiPhi29 Pol (0.5 U/µL). Aqueous phase is placed on ice. 200 µL of emulsifier is added to the 50 µL aqueous phase and vortexed for 5 min with Vortex-Genie 2 to generate emulsion. Emulsion is aliquoted into PCR strip and incubated at 42°C for 4 hours, 65°C for 15 mins, followed by 4°C hold. Emulsion is consolidated from PCR strips into 2 mL DNA LoBind tube, followed by addition of 1 mL Ethyl Acetate, 400 µL 1X TE, and 40 µL 3M Sodium Acetate. Emulsion is broken by vortexing components for 5 seconds then centrifuging at 20K RCF for 2 mins. Aqueous phase is aspirated from bottom of tube after centrifugation, and transferred to new 2 mL DNA LoBind tube. 1 mL of ice-cold EtOH is added, followed by short vortex for <5 secs or until white string-like precipitate is observed. Ethanol precipitated concatemers are spun at 18K RCF at 4°C for 10 mins, supernatant is gently discarded, and pellet is washed 2X with 70% EtOH, followed by elution in EB buffer (Circulomics). Ethanol precipitated high molecular weight (HMW) DNA is used for second strand synthesis in 40 µL final volume using the following components and concentrations: HMW DNA (less than 20 µL volume), 1X EquiPhi29 Buffer, dNTPs (1mM), DTT (1mM), Yeast Inorganic Phosphatase (0.025 U), BSA (0.5 mg/mL), EquiPhi29 Pol (0.5 U/µL). Template is initially self-annealed in the presence of Buffer and dNTPs only by incubating at 95°C for 3 mins, followed by snap cooling at 4°C for 3 mins. The additional EquiPhi29 components are then added after snap cooling. Reaction is incubated at 42°C for 4 hours, 65°C for 15 mins, followed by 4°C hold. After second strand synthesis, sample is Ethanol precipitated, and eluted in EB Buffer (Circulomics). HMW dsDNA is run on an 0.8% E-Gel NGS with a Lambda Ladder (HindIII digest). DNA larger than 22kb and is excised from gel and gel extracted using Zymo Clean Large Fragment recovery kit. HMW DNA is further size selected with Short-Read Eliminator kit from Circulomics and eluted in 30 µL EB.

### Nanopore Sequencing

Samples were library prepped using the Ligation Sequencing Kit (SQK-LSK109). All samples sequenced with MinION R9.4.1 flowcells, basecalled with Guppy basecaller 3.6.0. Quality Control is performed on reads using NanoPlot package (github.com/wdecoster/NanoPlot)

### Bioinformatics

Reads are basecalled with Guppy basecaller 3.6.0 using dna_*r9*.*4*.*1_450bps_hac*.*cfg* configuration file. Basecalled reads are concatenated into a combined fastq files using *cat* command. Long concatemers are split based on presence of MrHAMER hairpin using the Porechop package v0.2.4 (github.com/rrwick/Porechop) with options *--extra_middle_trim_bad side 0 --extra_middle_trim_good side 0* and with custom adapters.py containing the following sequences:

MrHAMER_Start: ATCTCTCTCAACAACAAC

MrHAMER_End: AACGGAGGAGGAGGAAAAGAGAGAGA

Gal10-proF_Start: GGTGGTAATGCCATGTAATATG

Gal10-proF_End: CATATTACATGGCATTACCACC

2U5B4F_Start: AGTAGTGTGTGCCCGTCTGTTGTGTGACTC

2U5B4F_End: GAGTCACACAACAGACGGGCACACACTACT

Reads are then size filtered using Filtlong package v0.2.0 (https://github.com/rrwick/filtlong) with *-- min_length 4000* option. Size filtered reads are then demultiplexed into a folder containing individual fastq files, each containing repeats originating from a single read. This is done using a custom python2 script qfilesplitterV3.1.py and using minimum block number (-b) of 8 or 10 to ensure sufficient error correction (the - b setting is a direct determinant of MrHAMER error correction). This is followed by another custom python3 script protocolV3.3.py that runs parallel instances of the following processes:

Reference mapping of each individual fastq file in demultiplexed folder to the HIV reference (pSG3.1 strain) using minimap2 v2.17-r954-dirty (in map-ont mode) to generate an paf alignment file. Racon v1.4.3 [59] (github.com/isovic/racon) consensus correction package uses paf files and each fastq file to generate a polished assembly in fasta format. This polished assembly in fasta format is used as a reference for the Medaka polisher v1.0.1 (github.com/nanoporetech/medaka) which uses fastq files from demultiplexed folder and *r941_min_high_g360* model to further sequence correct reads.

The pipeline yields a single fasta file containing individual reconstructed genomes with high accuracy (usually 0.1-0.2% error rate) which can be used for downstream analyses for sequence co-variation or SNV enrichment. For reads with a known reference sequence, the *assess_assembly* command from Nanopore’s Pomoxis package (github.com/nanoporetech/pomoxis) was used to determine error-profile of reconstructed genomes. To calculate coverage evenness, reconstructed genomes can be mapped to reference and visualized with *Tablet[60]*, or sorted and indexed using *samtools depth* to create a coverage map.

### Template Switching Rate Calculation

Calculated at two steps, after RT-PCR, and for entire MrHAMER pipeline. For sequencing of RT-PCR products. Sequencing reads were mapped to HIV Reference using *minimap2*. A custom python2 script *TS_Count*.*py* was used to determine the number of reads that span both barcode regions and contain either barcode, both barcodes, or no barcodes. Template switch rate is calculated by doing a ratio of recombinants (two barcodes or no barcodes) over the total number of reads spanning barcode regions. For MrHAMER template switching, the process is identical but uses reconstructed Gag-Pol genomes as an input.

### Detection of linked mutations using CoVaMa

MrHAMER outputs fasta files containing reconstructed genomes at high accuracy (using at least 8 repetitive units per genome). For downstream analysis, these reconstructed genomes are mapped to the HIV reference using *minimap2* and the *map-ont* preset. Co-Variation Mapper[17] v0.7 package is comprised of two python scripts. The first script is CoVaMa_Make_Matrices.py which is run with *minimap2* outputs with the following options *--Mode2 Nucs –SAM1 --PileUp_Fraction 0*.*005 NT*, and generates and populates matrices containing nucleotide contingency tables. Since our effective limit of detection of linked mutations is between 0.5 and 1%, we use a PileUp_Fraction of 0.005 so that only contingency tables with mutants present at a frequency >0.5% are considered (this is a function of both effective LOD for linked mutations and expected error rate). For the next step, matrices are analyzed using CoVaMa_Analyse_Matrices.py using outputs generated by previous steps with the following options *--Min_Coverage 5 -OutArray -Weighted NT*. Only contingency tables populated with more than 5 aligned reads are used, to allow for sufficient statistical power. Results are output in txt format with each row indicating a pair of mutations, along with their LD, weighted LD, R^2^, and contingency table values. The linked positions with LD or R2 values in the top 150 are then plotted in a bubble plot via R to visualize locations of co-variant mutations, along with their statistical significance.

### Antiretroviral therapy regimen and longitudinal sample viral loads

The patient’s initial treatment course included a nucleoside reverse transcriptase inhibitor (NRTI)-based regimen of lamivudine (3TC) and zidovudine (AZT), coupled with the non-nucleoside reverse transcriptase inhibitor (NNRTI) efavirenz (EFV) and protease inhibitor (PI) nelfinavir (NFV). Following an initial viral rebound 16 months after cART initiation, antiretroviral therapy regimen was changed twice, each time to a cocktail of three NRTIs which first included 3TC, AZT and abacavir (ABC), followed by 3TC, ABC, and Tenofovir (TDF). Both NRTI therapy changes failed to prevent to subsequent viral rebounds, with samples from the second NRTI-related viral rebound used for sequencing as a virological failure (VF) timepoint. Viral RNA was extracted from blood plasma, reverse transcribed, PCR amplified using our validated RT-PCR pipeline, and sequenced to high accuracy using MrHAMER. Each longitudinal sample was sequenced for 9-10 hours and yielded ∼1.7 Gb using the ONT MiniION R9.4.1 sequencing device **(Supplementary Table 3)**.

### Viral RNA extraction from patient samples

Frozen plasma aliquots from patients undergoing cART was obtained from UCSD CFAR (Davey Smith, MD). For each sample, 1mL plasma was transferred to 1.5 mL DNA LoBind tubes (Eppendorf) and samples were centrifuged for 75 minutes at 18,000g and 4°C to concentrate virus. After centrifugation, 860 µL of supernatant were carefully aspirated from top of tube and frozen at -80°C. Remaining 140 µL of concentrated virus was processed according to QIAamp Viral RNA Mini kit (QIAGEN) protocol, with only deviation from protocol being viral RNA elution in 30 µL of Nuclease free water. 11 µL of this eluate was used as input for subsequent RT and serial emulsion PCR as described before.

### Generation of sample-specific consensus sequence for use as reference for single molecule reconstruction

Given high HIV intra-host diversity (both at the SNV and indel level), a sample-specific reference is generated for each patient isolate prior to single molecule reconstruction [61]. Fastq reads generated after demultiplexing with qfilesplitterV3.1.py (each containing at least 10 repeating units) are concatenated, mapped to pSG3.1 reference, followed by racon and medaka sequence correction to generate a sample specific consensus assembly. For patient samples, this sample consensus assembly is used as a reference for subsequent single molecule error-correction using the protocolV3.3.py script.

### Generation of a patient-specific naïve reference using standardized HXB2 coordinates

For downstream analysis of error-corrected Gag-Pol reads, the Naïve consensus assembly for the patient sample is mapped to the HXB2 fasta reference (accession number K03455) using MAFFT v7.471 with the following options *--ep 20 --keeplength --addfragments*. This yields a Naïve reference sequence that preserves HXB2 ORFs and positional coordinates, while taking into account prevailing genetic background of the treatment naïve error-corrected Gag-Pol reads. Error-corrected single molecule reads, and haplotype sequences can then be mapped to this newly generated reference to determine patterns of single nucleotide variants (SNV) and Insertion/Deletion events.

### Generation of viral haplotype clusters from high accuracy Gag-Pol reads

High accuracy Gag-Pol reads from Naïve and VF samples are separately mapped to their respective sample-specific consensus sequence using minimap2 with options *-ax map-ont*. The resulting Naïve and VF SAM files are then used as input for CliqueSNV v1.5.4 with the following options *-m snv-pacbio -rn -fdf extended4 -tf 0*.*001*. The resulting outputs delineate clusters (haplotypes) of Gag-Pol reads that share mutation patterns, and includes FASTA sequence and percent enrichment of each haplotype cluster.

### Phylogenetic Analysis

Haplotype sequences obtained via CliqueSNV for Naïve and VF samples are concatenated along with Naïve consensus assembly containing HXB2 coordinates. A multiple-sequence alignment (MSA) is performed using Clustal Omega v1.2.1, with resulting alignments. Phylogenetic analysis is performed with ClustalW v2.1 using Neighbor Joining (NJ) clustering algorithm and 1,000 bootstraps. The resulting PHB file is then visualized with iTOL v5 with branch lengths proportional to genetic distance and tree rooted to treatment naïve reference.

### Waffle Plot analysis (LiME + Waffle plots)

To better understand the possible evolution path taken by the viral population under antiretroviral therapies, a custom python script, Linked-Mutation Extractor (LiME), is used to acquire the higher-order linkage information from each Gag-Pol genome. LiME is designed to classify long sequences into several groups basing on their unique combination of nucleotides at a series of positions (Genetic Patterns).

The major inputs of LiME are a BAM file containing the aligned target sequences and a list of positions. In order to accelerate the analyzing process, the list of positions given is firstly chucked into several smaller subgroups with 5 or less than 5 positions. Genetic pattern identification and read classification are done independently in each subgroup, after which all genetic patterns belonging to the same read are collected and combined, forming an entire pattern covering all the positions given. LiME generates a waffle plot using the PyWaffle python package and stores the analyzing results in a CSV file, which contains the detailed information of genetic patterns, the number and ID of the reads harboring each genetic pattern.

### Generation of protein translations to visualize insertions proximal to Gag cleavage sites

CliqueSNV Naïve and VF haplotypes are concatenated to single FASTA file. GeneCutter (https://www.hiv.lanl.gov/content/sequence/GENE_CUTTER/cutter.html) is used to generate codon alignments to the Gag region. Protein alignments are exported in FASTA format and imported to JalView v2.11.1.3 for visualization.

## Supplementary Figures

**Supplementary Figure 1.**
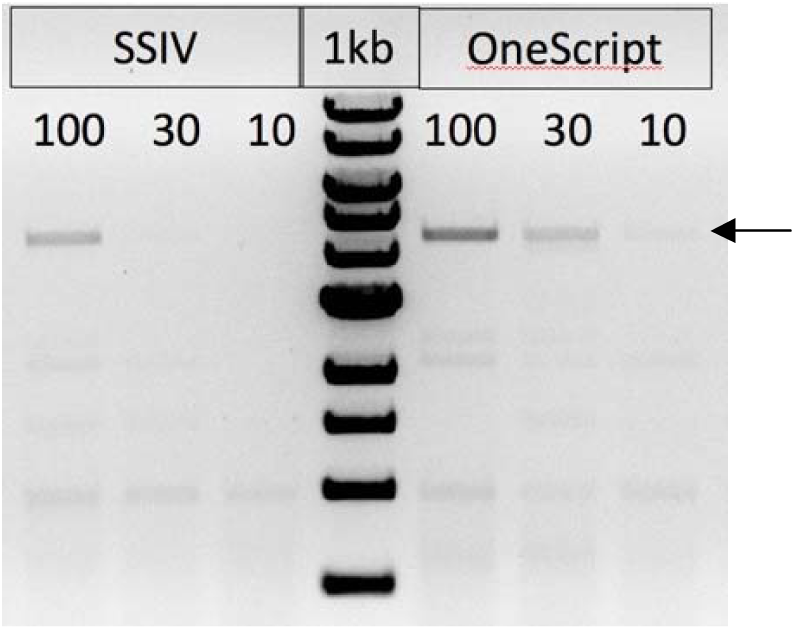
Priming Strategy and Determination of sensitivity of reverse transcription under limiting amounts of RNA inputs. A gene-specific primer (4609bp) was designed using the LANL HIValign DB to target the highly-conserved region proximal to the vif splicing junction in HIV RNA containing a flanking adapter (i.e. synthetic priming site) to be used for downstream PCR amplification. This priming strategy generates a ∼4.6kb cDNA product (arrow) that covers the Gag-Pol genomic region. This primer was used to reverse transcribe 100,000, 30,000 and 10,000 HIV RNA genome copies followed by PCR amplification for yield evaluation. Agarose gel electrophoresis of PCR amplification of cDNA obtained from Reverse Transcription using SuperScriptIV (SSIV) and OneScript Plus RT with OneScript Plus showing superior sensitivity compared to SSIV in producing cDNA from limiting amounts of input RNA.

**Supplementary Figure 2.**
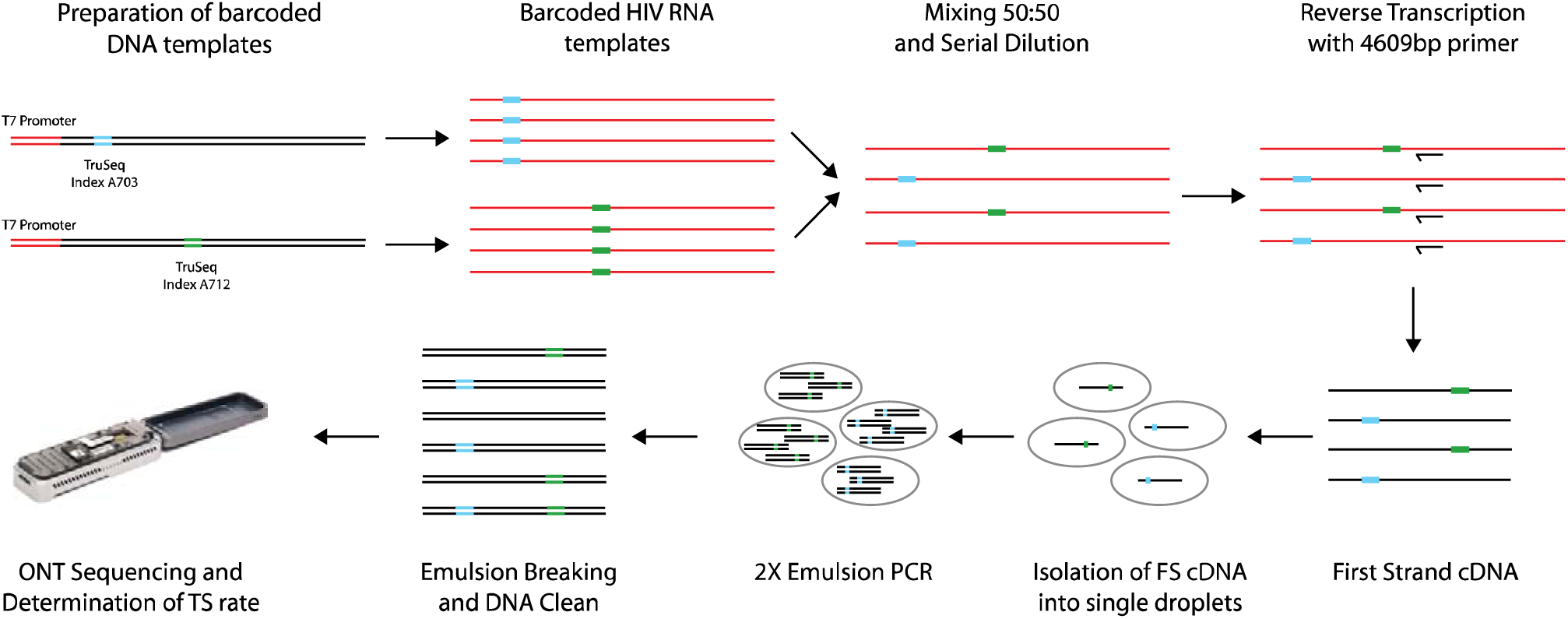
Nanopore based assay for direct readout of template switching for entire RT-PCR workflow. (A) An equal mixture of identical IVT HIV RNA templates, each containing a different 8-bp barcodes at 5’ and 3’ end, are serially diluted to 200K copies and used as input for reverse transcription, cDNA is PCR amplified, and Nanopore sequenced to determine incidence of cDNAs containing both barcodes or no barcodes, a direct indication of template switching.

**Supplementary Figure 3.**
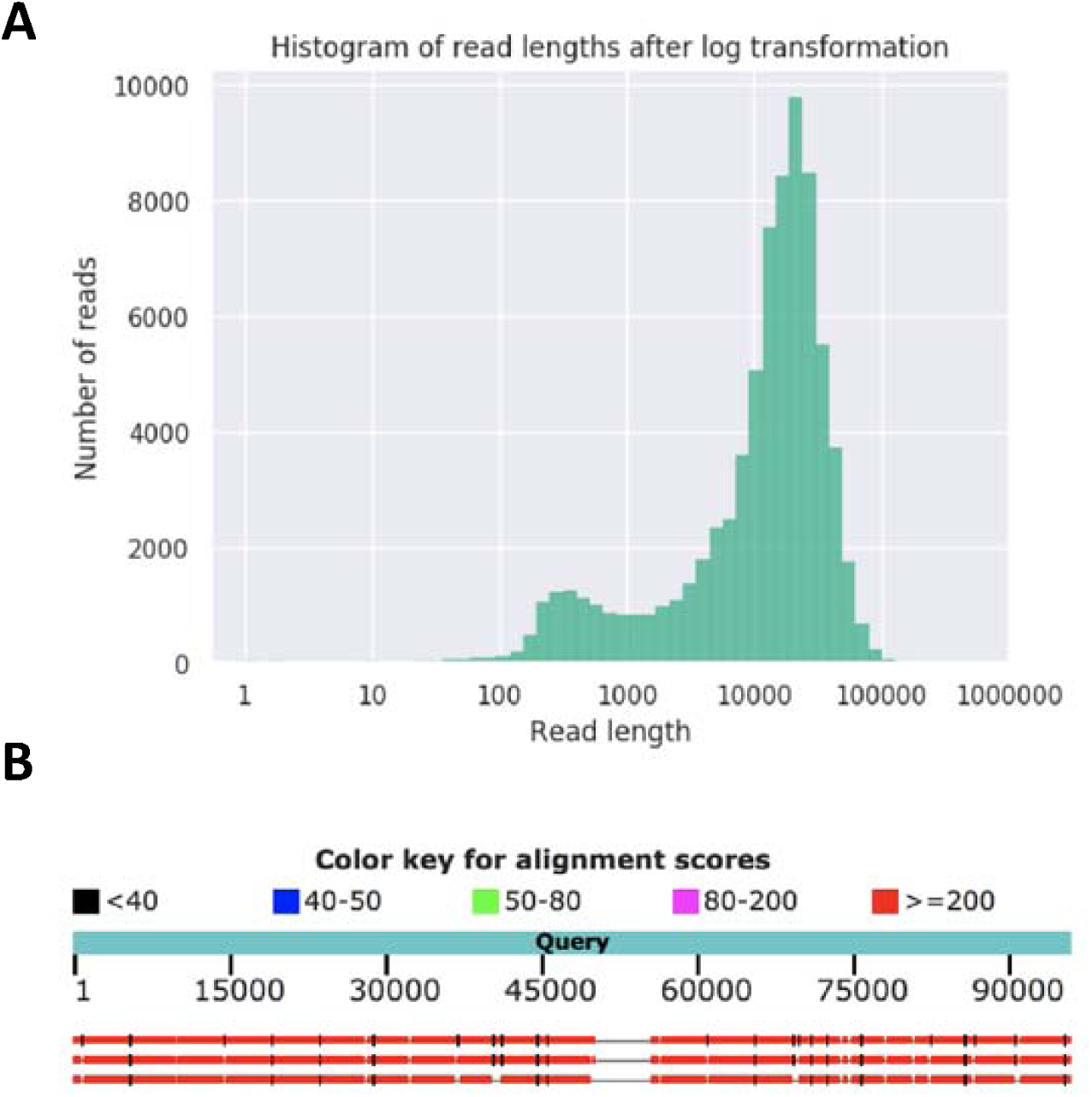
MrHAMER generates long concatemers that are detected via ONT sequencing. (A) Read length distribution of a typical MrHAMER sequencing run. Long read lengths with N50 values of ∼20kb are typical. (B) BLAST alignment of a 100kb read shows sequential repetitions of HIV genomes as expected.

**Supplementary Figure 4.**
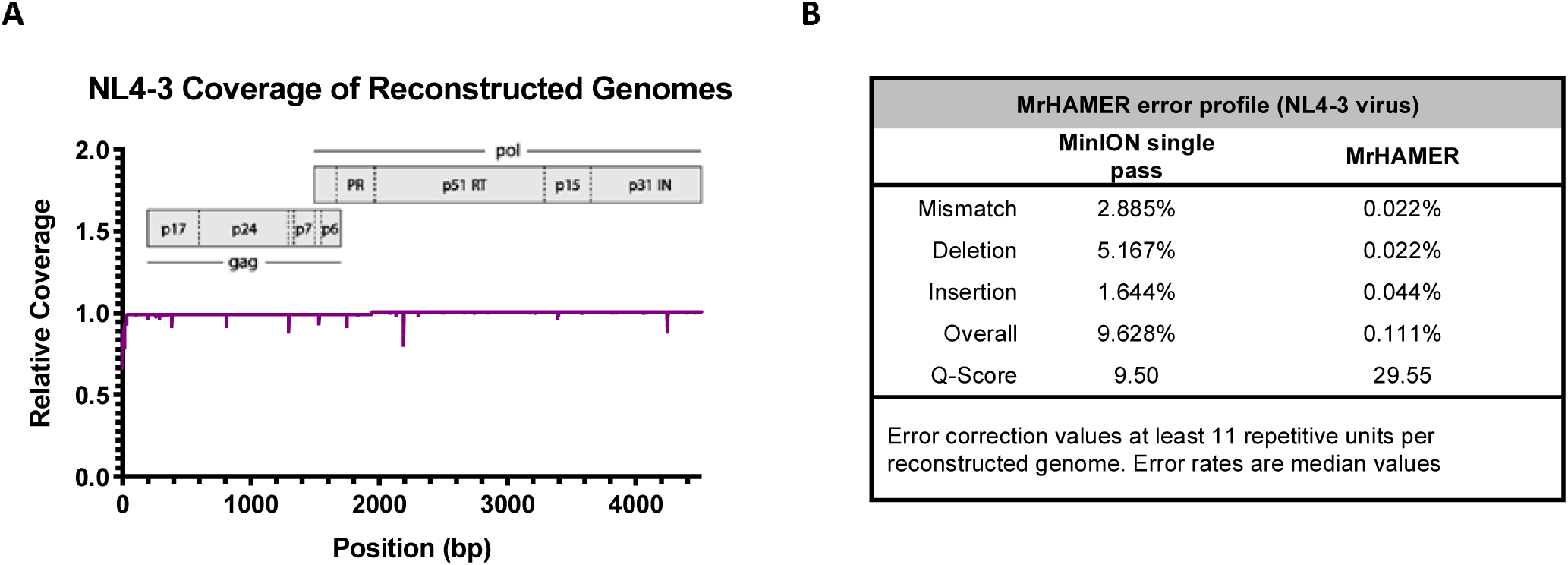
Full RT-PCR and MrHAMER pipeline validation using live NL4-3 virus generates high accuracy genomes with even coverage across Gag-Pol region. (A) Relative coverage of reconstructed NL4-3 Gag Pol genomes (B) Error profile of MrHAMER pipeline using NL4-3 viral inputs, quality scores approach Q30.

**Supplementary Figure 5.**
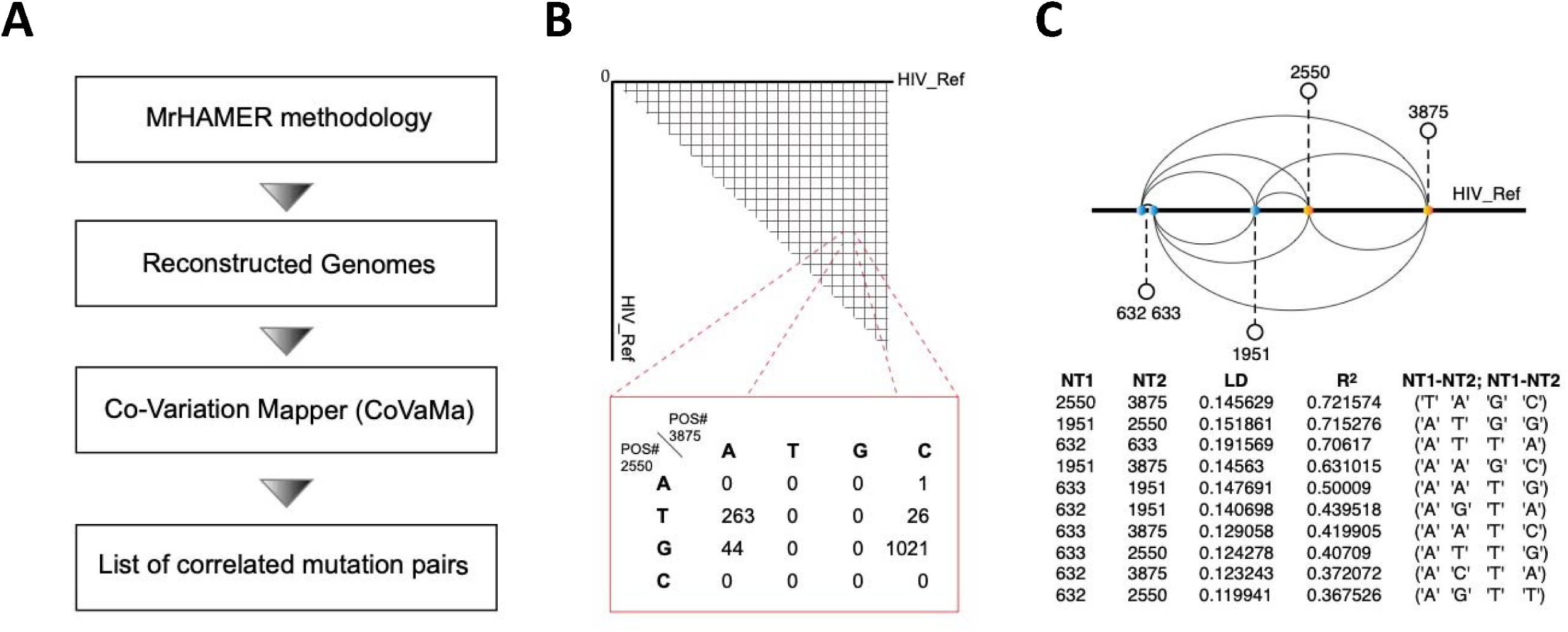
Workflow for analysis of pairwise nucleotide co-variation within genomes reconstructed via MrHAMER using Co-Variation Mapper (CoVaMa) (A) MrHAMER outputs are mapped to the HIV reference, and sequence alignment map outputs are used for CoVaMa analysis via two scripts (B) The first script evaluates every possible pairwise interaction within each genome via contingency tables, this is followed by the (C) Determination of highly significant interactions via linkage disequilibrium analysis and determination of R^2^ values.

## Supplementary Tables

**Supplementary Table 1.** Blood plasma obtained from the patient undergoing virological failure before treatment and after therapy failure.

| Specimen | PID | VIDate | Type | Volume (ml) | Viral Load | Treatment history |  |  |
| --- | --- | --- | --- | --- | --- | --- | --- | --- |
|  |  |  |  |  |  | Drug | Start_time | End_time |
| Treatment Naïve sample | 222-3jq | 11/10/1998 | Plasma | 1.0 | 208531 | None | None | None |
| Virological Failure sample | 222-3jq | 6/27/2003 | Plasma | 1.0 | 42000 | NFV/EFV/3TC/AZT | 01/13/1999 | 01/05/2001 |
|  |  |  |  |  |  | 3TC/AZT/ABC | 07/11/2002 | 07/20/2002 |
|  |  |  |  |  |  | 3TC/ABC/TDF | 02/08/2003 | None |

**Supplementary Table 2.** Genotypic drug resistance testing prior to initiation of cART showed no major drug-resistance mutants (DRMs) were detected.

|  |  |  |  |  |  |
| --- | --- | --- | --- | --- | --- |
| PID: 222-3jq |  |  | Specimen: Treatment Naïve sample | Type: plasma | Time: 11/18/1998 |
| Region | Start | End | Major mutations | Accessory mutations | Other mutations |
| PR | 1 | 99 | None | None | E35D,N37D,P39Q,D60E,L63P,I85IV |
| RT | 1 | 305 | None | None | K122E,T200A,R211K,V245A,S251I,V292I, E297A |
| IN | NA | NA | NA | NA | NA |

**Supplementary Table 3.** Sequencing information and statistics summary of the sequencing outputs for the Treatment Naïve sample (Left) and the Virological Failure sample (Right).

| Sample Information |  |
| --- | --- |
| PID | 222-3jq |
| Specimen | Treatment Naïve sample |
| Run Information |  |
| Flow cell type | FLO-MIN106 |
| Kit | SQK-LSK109 |
| Total run length (hours) | 10.5 |
| Statistics Summary |  |
| Mean read length | 15,916.6 |
| Mean read quality | 8.9 |
| Median read length | 13,015.0 |
| Median read quality | 8.8 |
| Number of reads | 97,755.0 |
| Read length N50 | 22,836.0 |
| Total bases | 1,555,924,055.0 |
| Number, percentage and megabases of reads above quality cutoffs |  |
| >Q5 | 90740 (92.8%) 1502.6Mb |
| >Q7 | 77942 (79.7%) 1299.7Mb |
| >Q10 | 26470 (27.1%) 332.5Mb |
| >Q12 | 9807 (10.0%) 72.9Mb |
| >Q15 | 2168 (2.2%) 8.0Mb |
| PID | 222-3jq |
| Specimen | Virological Failure sample |
| Run Information |  |
| Flow cell type | FLO-MIN106 |
| Kit | SQK-LSK109 |
| Total run length (hours) | 9.2 |
| Statistics Summary |  |
| Mean read length: | 15,249.4 |
| Mean read quality: | 8.9 |
| Median read length: | 12,386.5 |
| Median read quality: | 8.8 |
| Number of reads: | 116,248.0 |
| Read length N50: | 21,822.0 |
| Total bases: | 1,772,710,054.0 |
| Number, percentage and megabases of reads above quality cutoffs |  |
| >Q5 | 108184 (93.1%) 1712.9Mb |
| >Q7 | 93425 (80.4%) 1486.2Mb |
| >Q10 | 28862 (24.8%) 313.3Mb |
| >Q12 | 12088 (10.4%) 65.7Mb |
| >Q15 | 3725 (3.2%) 12.6Mb |

**Supplementary Table 4.** Positions of significant CoVaMa hits, associated LD and R2 values, and nucleotide identity of two most enriched contingency tables for given position

| NT1 | NT2 | wLD | wR2 | NT1 - NT2 | Nt1 - NT2 |
| --- | --- | --- | --- | --- | --- |
| 2550 | 3875 | 0.145629 | 0.721574 | T - A | G - C |
| 1951 | 2550 | 0.151861 | 0.715276 | A - T | G - G |
| 632 | 633 | 0.191569 | 0.70617 | A - T | T - A |
| 1951 | 3875 | 0.14563 | 0.631015 | A - A | G - C |
| 633 | 1951 | 0.147691 | 0.50009 | A - A | T - G |
| 632 | 1951 | 0.140698 | 0.439518 | A - G | T - A |
| 633 | 3875 | 0.129058 | 0.419905 | A - A | T - C |
| 633 | 2550 | 0.124278 | 0.40709 | A - T | T - G |
| 632 | 3875 | 0.123243 | 0.372072 | A - C | T - A |
| 632 | 2550 | 0.119941 | 0.367526 | A - G | T - T |

